# AlphaViz: Visualization and validation of critical proteomics data directly at the raw data level

**DOI:** 10.1101/2022.07.12.499676

**Authors:** Eugenia Voytik, Patricia Skowronek, Wen-Feng Zeng, Maria C. Tanzer, Andreas-David Brunner, Marvin Thielert, Maximilian T. Strauss, Sander Willems, Matthias Mann

## Abstract

Although current mass spectrometry (MS)-based proteomics identifies and quantifies thousands of proteins and (modified) peptides, only a minority of them are subjected to in-depth downstream analysis. With the advent of automated processing workflows, biologically or clinically important results within a study are rarely validated by visualization of the underlying raw information. Current tools are often not integrated into the overall analysis nor readily extendable with new approaches. To remedy this, we developed AlphaViz, an open-source Python package to superimpose output from common analysis workflows on the raw data for easy visualization and validation of protein and peptide identifications. AlphaViz takes advantage of recent breakthroughs in the deep learning-assisted prediction of experimental peptide properties to allow manual assessment of the expected versus measured peptide result. We focused on the visualization of the 4-dimensional data cuboid provided by Bruker TimsTOF instruments, where the ion mobility dimension, besides intensity and retention time, can be predicted and used for verification. We illustrate how AlphaViz can quickly validate or invalidate peptide identifications regardless of the score given to them by automated workflows. Furthermore, we provide a ‘predict mode’ that can locate peptides present in the raw data but not reported by the search engine. This is illustrated the recovery of missing values from experimental replicates. Applied to phosphoproteomics, we show how key signaling nodes can be validated to enhance confidence for downstream interpretation or follow-up experiments. AlphaViz follows standards for open-source software development and features an easy-to-install graphical user interface for end-users and a modular Python package for bioinformaticians. Validation of critical proteomics results should now become a standard feature in MS-based proteomics.

## INTRODUCTION

Mass spectrometry (MS)-based proteomics has evolved into a powerful and widely used analytical technique for researchers in diverse biological and clinical fields (1, 2). The increased throughput of MS instruments has led to the identification and quantification of thousands of proteins and their (modified) peptides in many experimental settings. To ensure the quality of such experiments, many journals now require to follow specific guidelines prior to submission (3). However, automated analysis workflows typically present long lists of identified and quantified peptides and proteins used for downstream analysis by the investigator. Only a small subset, like key proteins of signaling pathways and biomarker candidates are chosen for biological follow-up experiments or additional validation by orthogonal assays. Unfortunately, the underlying raw data for these critical peptides or proteins are rarely assessed at all or only in few of several possible dimensions, which could prevent investigators from following up on the best study candidates.

Applying the famous proverb “One picture is worth ten thousand words” to proteomics, visualization may be the most obvious solution for validating identifications at the level of raw MS data (4, 5). Inspecting the actual spectra of particular peptides, such as those with post-translational modifications (PTMs) or those uniquely identifying a protein of interest, can reveal important information, in addition to that used by the search engine. Furthermore, the advent of ultra-high sensitivity LC-MS based workflows for the analysis of minute protein amounts down to the level of single cells is currently lacking raw data visualization tools for the inspection and validation of proteins of interest at the limit of detection (6–9). The identification of a peptide amino acid sequence is part and parcel of high-confidence spectral identification and traditionally entailed visual inspection and validation by the investigator. However, the ever-increasing acquisition speed of mass spectrometers and the complexity of state-of-the-art scan modes in large-scale proteomics experiments rendered this approach impractical when dealing with huge data sets.

In part, this is due to many challenges in proteomic data visualization. The visualization step usually ranks last in the development of scientific algorithms or the establishment of novel workflows for analyzing proteomics data. Visualization tools tend to become publicly available with a considerable delay after the publication of the main workflows (10, 11). Because of their closed nature, even recently published tools may rapidly become outdated if they fail to take advantage of current advances in visualization such as in interactive biological ‘big data’ visualization. In this regard, increasing established open source concepts can help to keep up with the rapid pace of computational developments, building on powerful collaborative packages, for instance those in the increasing popular Scientific Python environment (12–14). In our group, we have focused on the visualization of highly complex multi-dimensional data acquired on Bruker TimsTOF instruments, which includes the additional ion mobility dimension (15). The AlphaTims package, as well as the parallel OpenTIMS effort, allows ready access and visualization of raw data, which has not been practical due to the long accession times and absence of convenient data structures (16, 17).

A major development in MS-based proteomics in recent years has been the success of machine learning in predicting peptide properties including retention time, ion mobility and the intensities of fragments in the MS2 spectra (18, 19). As a result, all these properties could be used to validate the proposed peptide spectrum matches, and this has already been done for spectral intensities (20). We reasoned that combining data visualization with the benefits of deep learning predictions, such as fragment ion intensities, retention time or ion mobility predictions, could dramatically benefit the entire visualization and validation approach. As a particular example, the assignment of convoluted fragmentation patterns in Data Independent Acquisition (DIA) to peptide sequences is still an active area of research with major search engines such as DIA-NN or Spectronaut sometimes disagreeing on the identification or matching of particular peptides (21, 22). Clearly, visualization of co-eluting fragments in the context of predicted retention times and fragment intensities (‘in silico truths’) could help in establishing confidence in critical peptide identifications.

Many of the currently existing visualization tools are proprietary and integrated into MS data analysis software pipelines by the MS manufacturer, such as Compass DataAnalysis (Bruker Daltonics), Freestyle and Xcalibur (Thermo Fisher Scientific), Sciex OS (Sciex) or by independent providers, such as Spectronaut and Skyline (22, 23). There are also standalone tools for visualization such as the PRIDE Inspector (24). However, in all these cases, these tools are difficult to reuse, extend, or integrate into existing workflows for example for novel multi-dimensional data, such as TIMS-TOF data.

Here, we developed a visualization tool with the following goals: It should allow (1) intuitive visualization of search engine results of the underlying raw data; (2) integration of *in silico* predictions by deep learning algorithms; (3) automation for end users through a graphical user interface or Jupyter Notebooks; (4) open-source accessibility and easy extendibility by bioinformaticians to incorporate new developments, for example interactivity, big data visualization and graph customization.

With these goals in mind, we developed AlphaViz, an open-source Python-based visualization tool that allows the user to explore identification and quantification confidence of peptides by visually comparing them to the signal presented in the unprocessed MS data. AlphaViz links identifications to the evidences of the raw data to assess their quality by using results from currently supported software tools, such as MaxQuant, AlphaPept and DIA-NN (10, 13, 21). It makes use of current advances in visualization, such as interactivity, “big data” visualization or real-time graph customization. The interactive plots included in AlphaViz provide the data on-demand in order not to overwhelm users, and include, for instance, zooming, selection and annotation. “Big data” capabilities make it possible to visualize millions of data points in a single graph in a browser. This enabled the visualization of MS heatmaps in AlphaViz, allowing to plot intensity of observed precursor masses across retention time and to visually assess MS peptide features in an enlarged view. In addition, customization of the plots, such as selection of a chart color scale or the size and format of the exported plots, enables researchers to easily create and extract illustrations of candidate proteins and peptides that are suitable for publication.

AlphaViz follows robust software development standards (high-quality code, extensive documentation, automated testing, and continuous integration) as a part of the AlphaPept ‘ecosystem’ (13).

## EXPERIMENTAL PROCEDURES

### Publicly Available MS Datasets

To demonstrate the use of AlphaViz for DDA data, we obtained raw data files of a fractionated HeLa library (fraction 1) generated with the 120-min gradient dda-PASEF method together with output of the MaxQuant software (v.1.6.1.13) from ProteomeXchange (data set PXD010012) (25). For the visualization of DIA data, we used a dataset previously acquired in our group: a 21-min gradient (60 samples per day) HeLa sample acquired on Evosep / timsTOF with the dia-PASEF method (data set PXD017703) (26). The results of DIA-NN analysis (v.1.7.15) of these data were taken from reference (27).

Additionally, we are presenting in-detail phosphoproteomics analyses. We used a recently published dataset where Hela cells were stimulated with EGF or left untreated, enriched for phosphopeptides and acquired in three replicates each on a timsTOF Pro instrument with a 21-min gradient and an optimal phosphoproteomics dia-PASEF method (28). The copied output of DIA-NN analysis (v.1.8) was filtered for 1 % PTM q-value, collapsed with the Perseus plug-in and filtered for 75 % localization probability (28).

### Data Acquisition for the Predict Mode Measurements

To demonstrate the ‘predict mode’ of AlphaViz, we synthesized phosphorylation positional isomers of the Rab10 peptide FHTITTSYYR. These isomers were dissolved in solution A* (0.1% TFA/2% ACN), and 125, 250, 500, 1250, 2500, and 5000 fmol of them were spiked into 50 fmol of bovine serum albumin. We measured the samples using a dia-PASEF method optimized for phosphoproteomics and 21 minutes Evosep gradients (60 samples per day method) combined with the timsTOF Pro (Bruker Daltonics) (28). The peptides were separated using an 8 cm x 150 μm reverse-phase column packed with 1.5 μm C_18_-beads (Pepsep) connected to a 10 μm ID nano-electrospray emitter (Bruker Daltonics). Our dia-PASEF method covered an *m/z*-range from 400 to 1400 Da and an ion mobility range from 0.6 to 1.5 Vs cm^-2^ with 12 dia-PASEF scans (cycle time: 1.38s). The collision energy depended on the ion mobility and changed from 60 eV at 1.5 Vs cm^-2^ to 54 eV at 1.17 Vs cm^-2^ to 25 eV at 0.85 Vs cm^-2^, and to 20 eV at 0.6 Vs cm^-2^.

### Design and Implementation

AlphaViz is written in Python, and its source code is freely available on GitHub (https://github.com/MannLabs/alphaviz) under the Apache license. The AlphaViz implementation combines a comfortable, reproducible and transparent working environment (Jupyter notebooks, GitHub, Binder, pytest) with a Python scientific stack consisting of highly optimized packages with elaborate testing, documentation and maintenance, allowing a focus on domain knowledge rather than implementation details (Fig. 1A). For data analysis in Python, we use NumPy for array manipulation, Pandas to handle tabular data, and Numba to speed up code execution with just-in-time code compilation. Furthermore, we use several open-source Python libraries for proteomics data analysis, such as AlphaTims to access Bruker ‘.d’ files and to convert them to Hierarchical Data Format (HDF) for fast reuse (16), and Pyteomics to handle ‘.fasta’ files (29). A set of well-established plotting libraries was used to generate all plots and a graphical user interface (GUI): (1) Bokeh, Plotly and Holoviews were used to build different types of interactive visualizations; (2) Datashader for fast visualization of large data sets; (3) Panel to implement a fully stand-alone GUI.

**Fig. 1.**
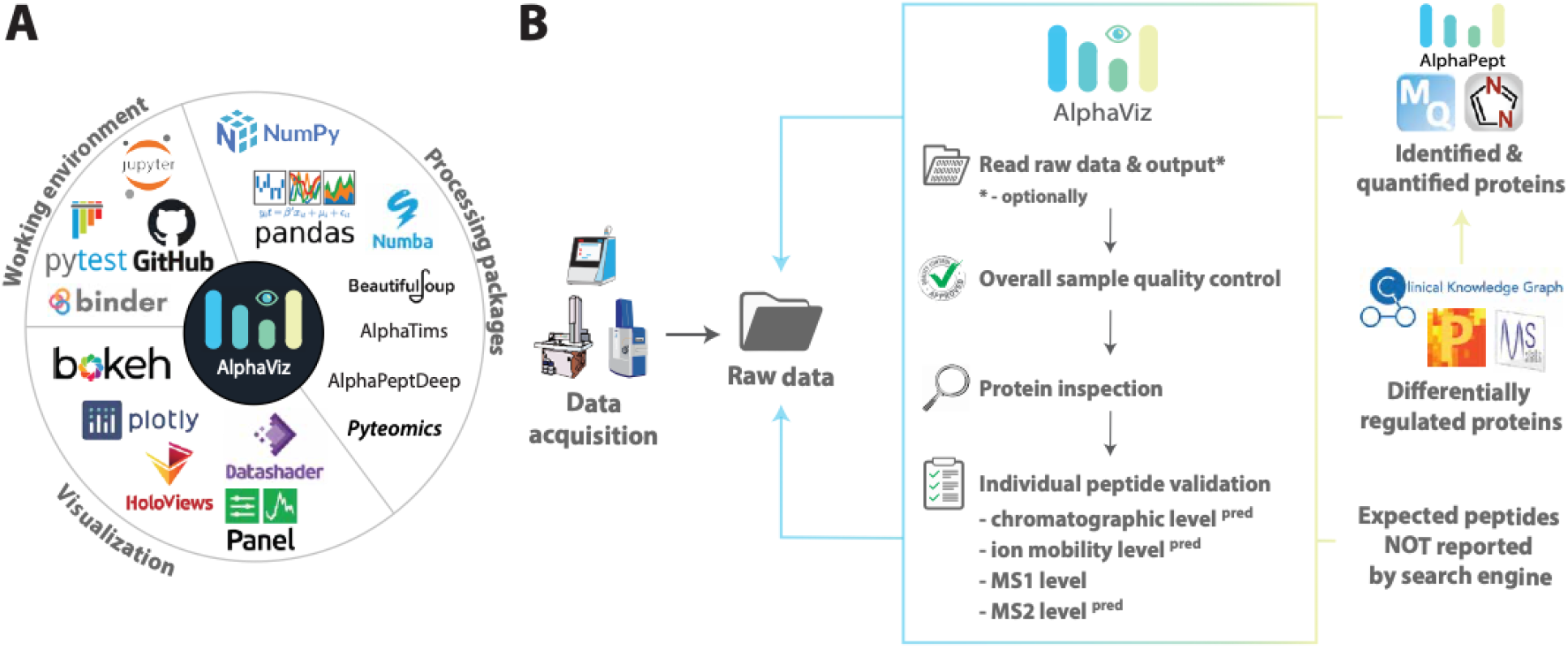
AlphaViz project dependencies and workflow. *A, Python libraries and other services used in AlphaViz*. The Python libraries and services fall into three groups, those for (1) efficient working and test environment; (2) data preprocessing and handling; and (3) visualization and the graphical user interface. *B, Overview of the AlphaViz workflow*. First AlphaViz directly reads the raw data together with the results of the supported proteomics workflows, reporting identified and quantified proteins of interests i.e. differentially regulated proteins. The overall sample quality can then be assessed using various quality metrics as a basis for further evaluation. Next, the user can inspect the individual quality of the critical proteins as well their identified peptides through AlphaViz. This is done at different levels, such as LC, IM, MS1 and MS2 levels, which can also be predicted using the built-in deep learning models for comparison. The ‘predict mode’ also allows to retrieve the signals from the raw data for peptides of interest that were not reported by the search engine (see Results for further explanation).

The AlphaViz implementation in the GitHub repository is organized into independent functional modules: (1) a ‘data’ folder with some necessary tables for performing calculations; (2) an ‘io’ module providing functionality for reading output files of proteomics data analysis programs; (3) a ‘preprocessing’ module that includes data preprocessing functionality; (4) a ‘plotting’ module containing all functions creating plots; (5) a ‘utils’ module including common utilities; (6) and a ‘gui’ module containing the entire implementation of the AlphaViz GUI. The helper units include: (1) a ‘style’ folder with files specifying the style of the dashboard elements; (2) an ‘img’ folder with logos and static images included in the GUI; (3) a ‘docs’ folder including a comprehensive GUI user guide. Besides the modular ‘alphaviz’ folder, the repository contains additional important information such as: (1) an ‘nbs’ folder with Jupyter Notebooks as tutorials for AlphaViz as a Python package usage; (2) the ‘test’ and ‘test_data’ folders containing functions which test the functionality of all previously mentioned Python modules and the necessary test data for them; (3) a general .README file with details on installation, usage of the different AlphaViz modes (GUI or a Python package), contributions and much more; (4) a ‘requirements’ folder with specific dependencies; (5) all other folders, e.g. ‘misc’, ‘docs’, ‘.github’ and ‘release’ that are involved in a continuous integration pipeline with automatic testing, creation of GUI installers for all OSs, the release of the new versions on GitHub, PyPI (https://pypi.org/project/alphaviz/) and ‘Read the Docs’ (https://alphaviz.readthedocs.io/en/latest/).

### Modes

Depending on users’ programming skills, AlphaViz can be operated in two modes: a user-friendly browser-based GUI and a well-documented and tested module with Python functionalities.

The AlphaViz GUI has one-click installers provided on the GitHub page for Windows, macOS and Linux (https://github.com/MannLabs/alphaviz#one-click-gui). A comprehensive AlphaViz user guide is provided on GitHub.

AlphaViz can be installed from PyPI using the standard pip module. Compared to the GUI, this mode provides more flexibility for users with programming experience, allowing reuse of the plotting or data importing or preprocessing functions to reproduce the same analysis and visualization. To facilitate the use of AlphaViz as a Python package and to lower the entry barrier for users, we created Jupyter notebook tutorials separately for the different available pipelines: for DDA data analyzed with MaxQuant, for DIA data analyzed with DIA-NN, and for the targeted mode without any prior identification. The tutorials offer code to reproduce the results obtained in the GUI.

### Quality metrics

We include the following statistical distributions of peptide data which should be checked in AlphaViz to ensure good data quality:

- for DDA data analyzed by MaxQuant (sixteen parameters): m/z, Charge, Length, Mass, 1/K0, CCS, K0 length, Missed cleavages, Andromeda score (peptide score), Intensity, Mass error [ppm], Mass error [Da], Uncalibrated mass error [ppm], Uncalibrated mass error [Da], Score (protein score), (EXP) # peptides (the number of experimentally found peptides);
- for DDA data analyzed by AlphaPept (eighteen parameters): m/z, Charge, Mass, IM, Length, delta_m, delta_m_ppm, fdr, prec_offset_ppm, prec_offset_raw, hits, hits_b, hits_y, n_fragments_matched, (EXP) # peptides, q_value, score, score_precursor;
- for DIA data analyzed by DIA-NN (seventeen parameters): m/z, Charge, Length, IM, CScore, Decoy.CScore, Decoy.Evidence, Evidence, Global.Q.Value, Q.Value, Quantity.Quality, Spectrum.Similarity, (EXP) # peptides, Global.PG.Q.Value, PG.Q.Value, PG.Quantity, Protein.Q.Value.

## RESULTS

We developed AlphaViz to visually validate critical proteins and peptides at the raw data level. It currently supports timsTOF data acquired in data-dependent acquisition (DDA) or data-independent acquisition (DIA) mode. As detailed in the Experimental Procedures, AlphaViz is written in Python using various open-source libraries for data accession, analysis and visualization. To ensure that the tool can be used by a wide audience, AlphaViz is available on Windows, macOS, and Linux, in two different modes: a convenient graphical user interface (GUI) and as a well-documented and tested Python package.

AlphaViz works either with the output of proteomics software pipelines or only at the raw data level. Using the output results, it first enables the overall quality of a particular sample to be assessed, as a basis for further automated analysis. It then superimposes the identifications provided by common proteomics workflows, such as MaxQuant, AlphaPept, or DIA-NN, on the raw data signals.

For integrating in silico predictions of experimental peptide predictions from the (modified) sequences, we use our AlphaPeptDeep package that itself is built on the pDeep model (30–32). In contrast to pDeep, AlphaPeptDeep supports not only MS2 prediction but also retention time (RT) and collisional cross section (CCS) prediction for any peptide modification (33, 34). Furthermore, AlphaPeptDeep provides easy-to-use transfer learning functionalities that were also used in AlphaViz to fine-tune the experiment-specific RT predictions.

These readily available *in silico* predictions for any peptide sequence, enables a distinct ‘predicted mode’ whereby the calculated coordinates of a peptide sequence of interest are projected onto the raw data. A variety of its applications are shown below.

In the following, we employ several use cases or examples to describe the entire validation procedure for peptides of several specific proteins using DDA and DIA data, pinpointing unreliable peptides although they were highly scored by software analysis tools. We then show applications of peptide signals retrieved directly from the raw data based on the predicted or experimental properties of the peptides. Finally, we illustrate the use of AlphaViz to explore critical nodes in the phosphoproteome of the EGF signaling pathway.

### Visual validation of global parameters and individual peptides of critical proteins

Today, researchers typically at best inspect a few examples of all detected peptides, yet rarely the main biological or clinical hits that are the main results of the project. Furthermore, the overall quality of the proteomics dataset is often not examined at the level of MS results. This is partly because it may not be easy in the available software to assess these crucial parameters and results, especially for dda-or dia-PASEF data. Clearly, it would be desirable to be able to confirm at a global level of each LC-MS run that the proteomics data is free of major issues or biases so there is a solid basis for further evaluation. After that, verification should be done individually for each protein of particular interest at the peptide level. This will help to increase confidence in the protein identifications reported by the search engines or result in discarding the identification during the various quality checks as illustrated below.

In Figure 2A we exemplify the entire validation process applied to the DDA data, which originates from a HeLa sample acquired on a timsTOF instrument with a 120-min gradient in dda-PASEF mode and analyzed by MaxQuant (Fig. 2A, Experimental Procedures) (10). We first imported all raw data and MaxQuant results. AlphaViz then displays the overall quality metrics of the raw data to ensure the quality of the MS runs, which is shown for Fraction 1 as an example (Fig. 2B). In the total ion chromatogram (TIC) and base peak intensity (BPI) chromatogram the typical shape and overall high stable intensity level of the MS1 and MS2 TIC reveal no anomalies (Fig. 2B). The MS1 and MS2 BPI also indicate no major issues with saturation of the LC-MS system, such as overloading or contamination (Supplementary Fig. S1A). To dig deeper into the raw data quality, we suggest using AlphaTims, which quickly displays any desired slice of the billions of raw data points (16). Next, we selected six metrics available in AlphaViz to obtain an overview of all the peptides identified by MaxQuant, which revealed typical distributions for m/z values, peptide lengths, ion mobility values, and number of peptides per protein (Fig. 2B, Supplementary Fig. S1B, Experimental Procedures). However, a clear overall mass-shift is apparent. This is caused by AlphaViz using the raw data directly instead of re-calibrated values after a first database search. When inspecting individual peptides (see below), the re-calibrated mass measurements are used, with a user-definable tolerance, i.e. for visualizing extracted ion chromatogram (XIC) traces. Many peptide metrics are relative to an overall distribution and visualizing their position respective to the raw data with AlphaViz allows context-specific interpretation. For instance, the Andromeda score (MaxQuant) at 1% FDR shows an interquartile range (IQR) between 47 and 100 with 379 outliers above 177, suggesting that values above 100 should have a very high probability to be correct.

**Fig. 2.**
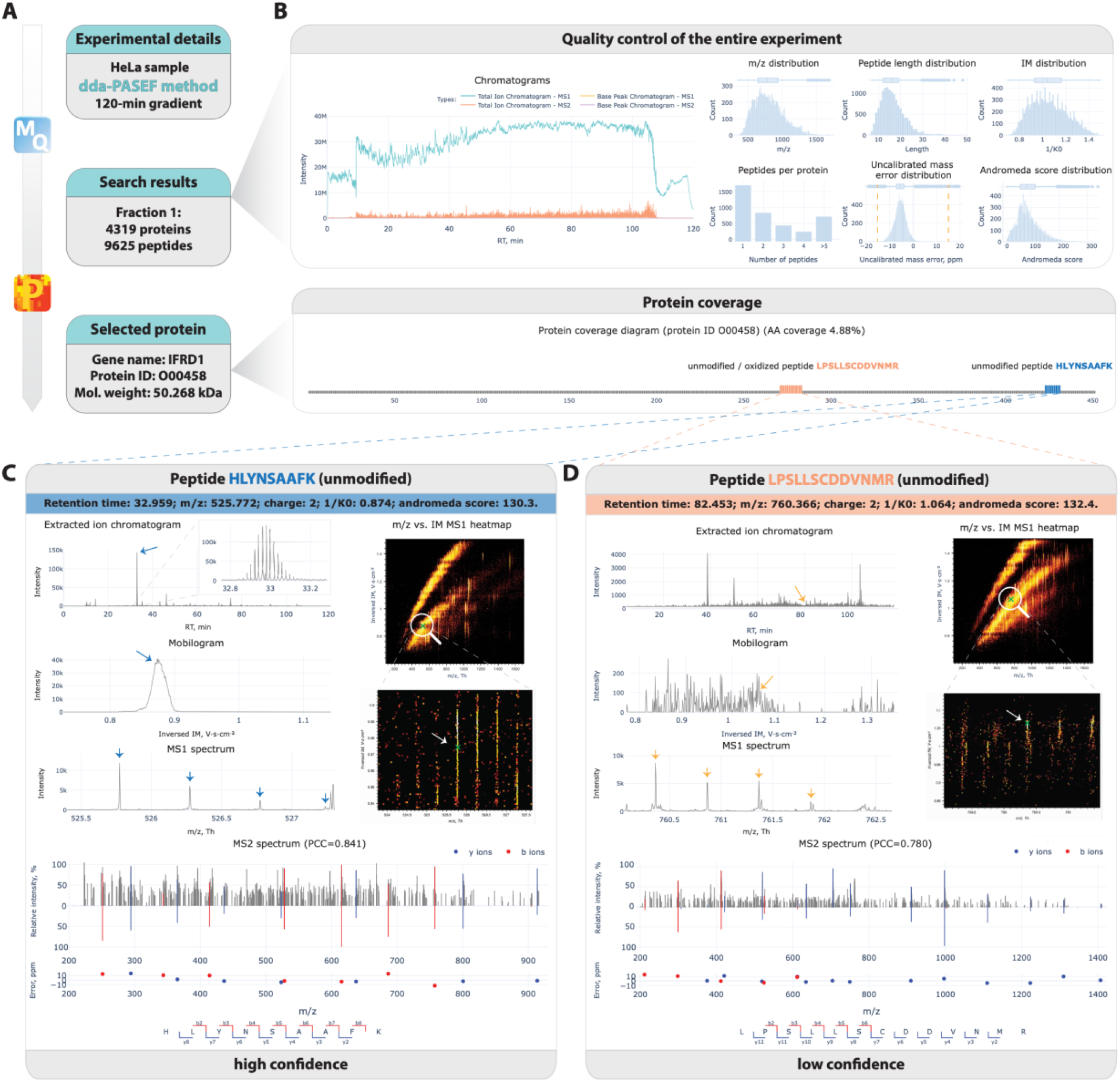
Validation pipeline in AlphaViz of two unmodified peptides of the same protein using timsTOF DDA data analyzed by MaxQuant. *A, Workflow*. A fractionated 120-min HeLa sample, acquired with dda-PASEF and analyzed by MaxQuant (PXD010012) (25), was imported in AlphaViz. *B, Overall sample quality*. Chromatograms and additional quality metrics of fraction 1. Interferon-related developmental regulator 1 protein (ID O00458) was selected for further detailed exploration. The “Protein coverage” bottom panel shows the identified peptides in the sequence context of the protein (similar to, but less detailed than AlphaMap (35)). *C and D*, The visualization of XIC, mobilogram, MS1 spectrum with overall and zoomed MS1 heatmaps together with the experimental and predicted MS2 spectrum. *C, Peptide view*. Inspection of the unmodified peptide HLYNSAAFK reveals it to be a high confidence identification. *D, Peptide view*. The unmodified peptide LPSLLSCDDVNMR has low confidence.

We next inspected peptides of interferon-related developmental regulator 1 protein (ID O00458) to represent a protein of particular biological importance that was identified with only a few peptides and a relatively low protein score of 35 which is in the first quartile of the distribution (Supplementary Fig. S1B). The protein q-value (probability to be wrongly identified) is only about 10^−4^, derived from the peptide posterior error probabilities (PEPs) of its two identified peptides, one of them also as an oxidized form. The Andromeda scores of the unmodified peptides are 130.3 and 132.4, well within the highest quartile with a PEP of less than 0.6%. Because of the discrepancy of these high peptide scores and the low protein score, we visualized the underlying raw peptide data in AlphaViz.

We first assessed the XIC of the unmodified peptide HLYNSAAFK (± 15 ppm, ± 0.05 1/K0, Fig. 2C). This revealed a pronounced peak at the reported retention time of 32.96 min, close to the value of 34.82 min predicted by AlphaPeptDeep. Moreover, the peak shape was Gaussian with limited tailing. Similarly, the extracted ion mobilogram (ppm and retention time window of ± 15 ppm and ± 30 seconds) shows a narrow peak at the reported 1/K0 of 0.874, almost identical to 0.892 predicted by AlphaPeptDeep. This also illustrates the advantage of the additional ion mobility dimension to evaluate the quality of peptide identifications.

AlphaViz can also visualize the MS1 context from which the precursor was picked for sequencing, in this case revealing a well-defined feature in the m/z and ion mobility dimensions (the entire heatmap with the zoomed view in Fig. 2C). All fragment ions for this particular peptide are present in the MS2 spectrum with an average absolute mass error of 3.1 ppm. The spectrum predicted by deep learning in the mirrored spectrum has a similar intensity pattern as the measured one (Pearson correlation coefficient of 0.841, Fig. 2C, bottom panel). Although some peaks remain unidentified, most of the larger peaks are correctly annotated.

We then examined the second unmodified peptide LPSLLSCDDVNMR. Despite its similarly high score, and PEP of 10^−4^, its XIC was two orders of magnitude less intense and without a well-defined peak shape at the claimed retention time (82.45 min; predicted 83.28 min). Similarly, the extracted ion mobilogram also lacks the expected clear peak shape at 1.064 1/K0 (predicted 1.038 1/K0, Fig. 2D). In comparison to the previously analyzed peptide, it is apparent in the heatmap for the MS1 frame that the peptide of interest was picked in a crowded region, which could potentially lead to a chimeric MS2 spectrum. This goes along with its fuzzy MS1 feature in the *m/z* and ion mobility dimensions and a relatively large mass deviation of around 50 ppm (Fig. 1D). In addition, the MS1 spectrum reveals an isotope pattern with some interference from another precursor. However, when inspecting the MS2 spectrum, many ions from the b- and y-series were identified by MaxQuant with a mean mass error of 0.3 ppm and demonstrated a similar intensity pattern with the predicted mirrored spectrum (Pearson correlation coefficient of 0.780, Fig. 2D, bottom panel). This turned out to be the reason for the high Andromeda score of the peptide, which is based on the number of detected fragment ions. Nevertheless, both the low values of the overall absolute peak intensities in the MS2 spectrum (below 300) and the poor data quality in other above-mentioned dimensions suggest that this peptide is a false positive hit despite the high peptide score. Thus, we illustrate the use of AlphaViz to evaluate two identified peptides of the same protein reported by MaxQuant with similar scores, only one of whom should be considered as a reliable hit according to our analysis.

### Visual validation of peptidoforms of the proteins of interest

In recent decades, MS has become the tool of choice for large-scale identification and quantitation of proteins and their post-translational modifications (PTMs) and the computational workflows for the analysis of DDA have matured. DIA analysis is comparatively newer and less established, especially when PTMs are being analyzed (28, 36–39). Although modern proteomics workflows report the localization of the identified PTMs and the associated probabilities, in our experience it is still necessary to manually validate the results for individual proteins and PTMs of critical importance. Compared to DDA data, validation of peptides at the raw level in the DIA pipeline should additionally include detailed inspection at the precursor retention time and, if applicable, ion mobility values. The values extracted from the library should then match the values in the raw data within the experimental error, especially the coelution of matched fragments and precursors.

Figure 3A presents the entire validation process of peptidoforms applied to a HeLa sample acquired on a timsTOF Pro instrument with a 21-min gradient in dia-PASEF mode and analyzed by DIA-NN (Experimental Procedures). We first imported the raw data file of the selected sample along with the DIA-NN output result into AlphaViz. The overall sample quality panel in AlphaViz was used to evaluate the overall quality of the selected sample (A5_1_2451) (Fig. 3B). The TICs and BPIs for MS1 and MS2 levels demonstrate typical shapes and overall high level of intensity without any visible anomalies (Fig. 3B, Supplementary Fig. S2A). As before, for further quality checks, such as verification of mass calibration or ion mobility stability, we suggest using AlphaTims (16). By selecting six out of seventeen available quality metrics, we observed typical distributions of important parameters, such as peptide *m/z*, ion mobility values and a preponderance of double-and triple-charged ions (Fig. 3B, Supplementary Fig. S2B, Experimental Procedures). Furthermore, a high number of peptides identifying each protein also serves as an important quality assurance. For our example, the peptide score distribution (Quantity.Quality score) in DIA-NN for each individual peptide at 1% FDR shows an IQR between 0.67 and 0.91, suggesting that scores above 0.91 (top 25% of the significant scores) should be correct with a high probability.

**Fig. 3.**
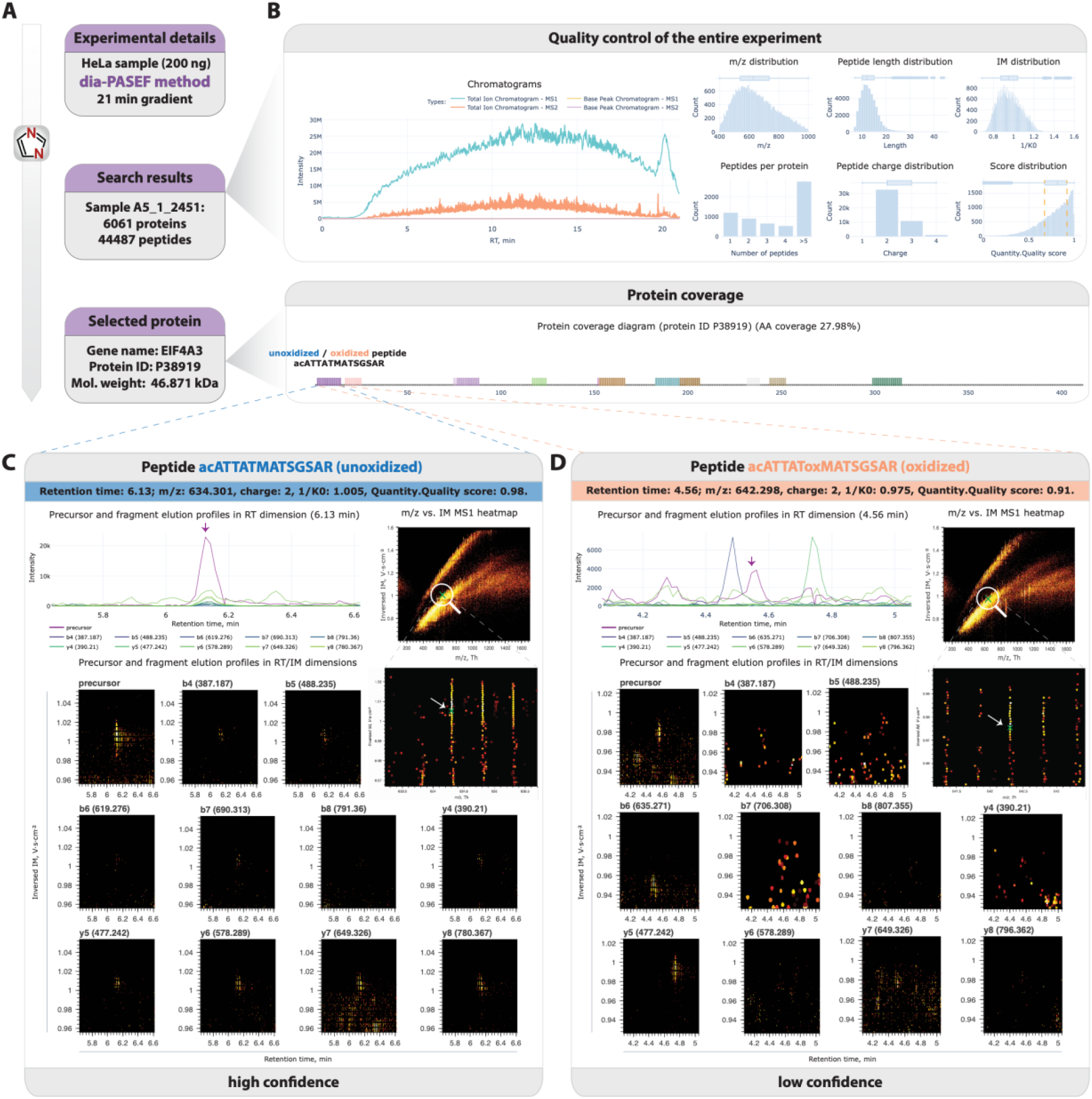
Validation pipeline in AlphaViz of two peptide variants using timsTOF DIA data analyzed by DIA-NN. *A, Workflow*. A 21-min HeLa sample, acquired with dia-PASEF and analyzed by DIA-NN (PXD017703), was imported into AlphaViz (26, 27). *B, Overall sample quality*. Chromatograms and additional quality metrics of fraction 1. Eukaryotic initiation factor 4A-III protein (ID P38919) was selected for further detailed exploration. The “Protein coverage” bottom panel shows the identified peptides in the sequence context of the protein. *C and D*, Visualization of overall and zoomed-in MS1 heatmaps, precursor and fragments elution profiles in both retention time (line plots) and retention time and ion mobility (heatmaps) dimensions. *C, Peptide view*. The unoxidized N-terminal acetylated peptide ATTATMATSGSAR shows high confidence. *D, Peptide view*. The oxidized peptidoform of the same peptide demonstrates low confidence.

Satisfied with the overall sample quality check, we chose eukaryotic initiation factor 4A-III protein (ID P38919, q-values < 10^−4^) for further investigation because of its high sequence and PTM coverage including a total of 20 peptide variants that mapped to thirteen peptides with q-values < 5*10^−3^. Two of these peptidoforms were the unoxidized and oxidized forms of the N-terminally acetylated peptide ATTATMATSGSAR, which were reported with similar scores by DIA-NN of 0.98 and 0.91 respectively.

To investigate if both forms were actually present, we first assessed the MS1 heatmap of the unoxidized peptide ATTATMATSGSAR (Fig. 3C). The extracted position of the peptide on the *m/z* versus ion mobility MS1 heatmap revealed a well resolved feature (Fig. 3C). The elution profiles of its precursor with all fragment ions (± 30 ppm, ± 0.05 1/K0, ± 30 sec) likewise demonstrated a sharp high-intensity precursor peak at 6.13 min (predicted 6.29 min), which coelutes with almost all of the main fragment ions. Taking advantage of ion mobility to ensure the presence of the MS signals, we also visualized the heatmaps for the precursor and each individual fragment in retention time and ion mobility dimensions colored by intensity. These heatmaps confirm the presence of the analyzed peptide.

Investigation of the MS1 heatmap of the oxidized form of the same peptide revealed a comparably well-defined MS1 feature (Fig. 3D). However, analysis of the elution profiles in the retention time dimension only showed a low-intensity precursor peak at 4.56 min (predicted 4.07 min) with only few and unaligned peaks. Conversely, the heatmaps for the precursor ion and its fragment ions confirm the absence of fragment ions signals within the expected retention time and ion mobility ranges. Correct identification of modified peptides in DIA data is a known challenge and is thought to be impeded by the presence of shared unmodified fragments of the base peptide in the DIA matching library (38).

### Validation of peptides directly in DIA raw data with the ‘predict mode’ in AlphaViz

The processing of DIA data is computationally challenging due to the high complexity of MS2 spectra containing fragments of multiple precursors from each single isolation window. Peptides with low signal intensities can be difficult to detect and easy to misinterpret. Conversely, quality measures of such peptides, provided as q-values, often fall below preset thresholds, resulting in “missing values”, even though the peptide is actually present. We hypothesized that by manually visualizing their signals in AlphaViz, they could still be extractable from the raw data. This is enabled by the integration of deep learning assisted prediction of retention time and ion mobility of AlphaPeptDeep in AlphaViz. We tested our hypothesis with two use cases: one regarding the detection of positional isoforms of a synthetic phosphopeptide and one regarding the retrieval of missing values.

We had previously identified the Rab10 protein as a clinically important substrate in Parkinson’s disease (40, 41). In the course of developing an assay to measure the phosphorylation site occupancy, we had synthesized positional phosphoisomers of the Rab10 peptide FHTITTSYYR (Fig. 4, left panel) (42). When analyzing these peptides by DIA software, they were not reported to be present (Experimental procedure). By predicting the retention time and ion mobility values for the different charges of two known phosphoisomers, their elution profiles were easily detected in the raw data of the two highest concentrations (Fig. 4, right panel, Supplementary Fig. S3). The XICs (+-30 ppm, +-0.05 1/K0, +-30 sec) demonstrate clearly defined high-intense precursor peaks with coeluting b3-b6 and y3-y6 fragment ions. The presence of these fragment ion signals is further confirmed by heatmaps that take advantage of the additional ion mobility dimension. Note that there is a slight difference between the predicted and actually observed retention time, which is not unexpected given the estimated accuracy of the prediction (32). Given the co-elution behavior of the expected fragments in the AlphaViz, we confirmed the presence of the intended phosphoisomers and concluded that they were not detected in all samples because of low MS signals.

**Fig. 4.**
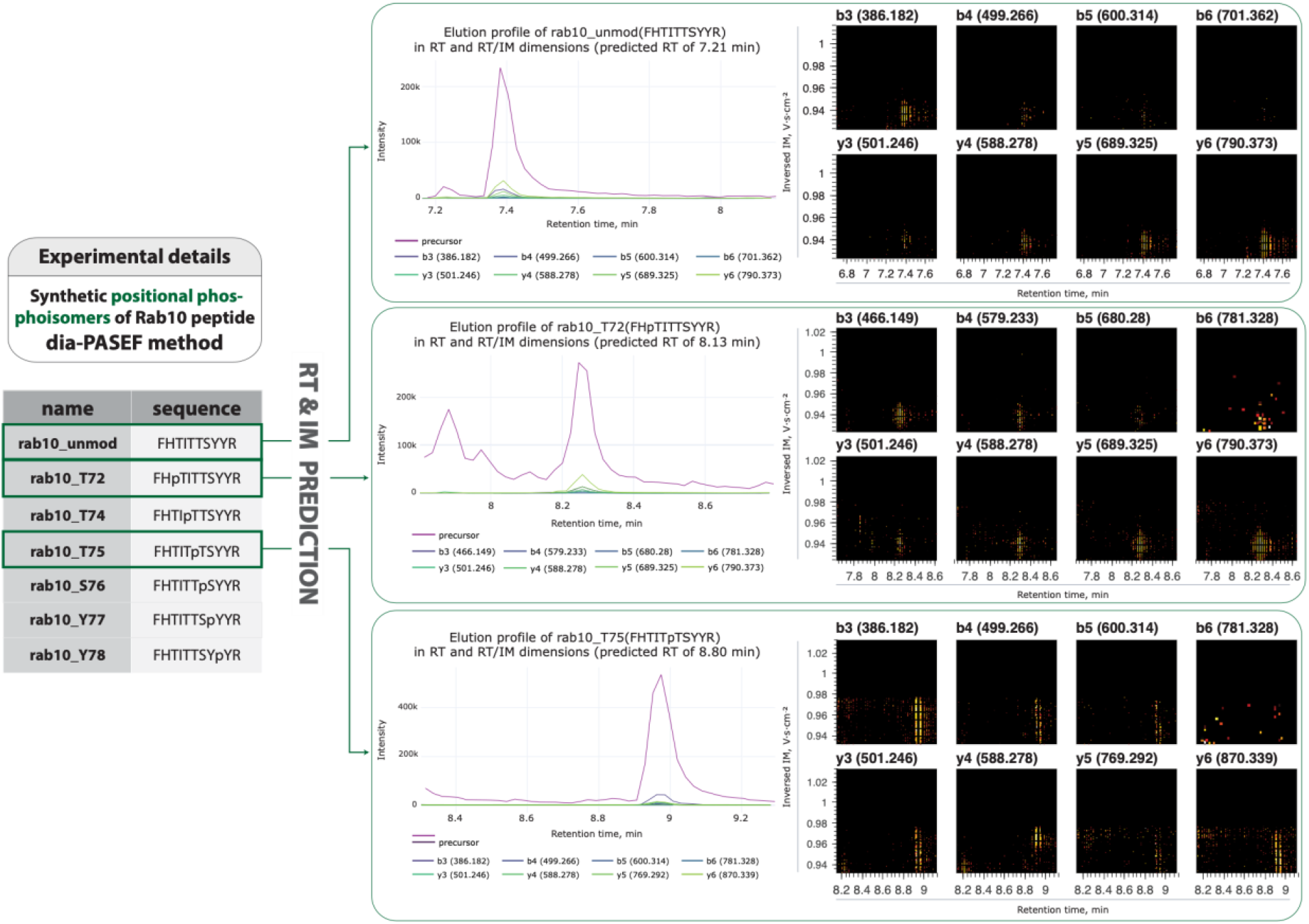
Validation of the presence of synthetic phosphoisomers of Rab10 peptide in DIA raw data. *Left panel*, Synthetic positional phospho-isomers of the Rab10 peptides. *Right panel, Extracted peptide signals for the sequences with a green box in the left panels*. Heat map of transitions from the raw data for the unmodified peptide and its two phosphoisomers.

To test if AlphaViz could retrieve seemingly missing values, we used the same HeLa DIA dataset as for the visual validation of peptide-forms above (Experimental Procedures). For illustration, we selected the cysteine-carboxylated peptide LCYVALDFEQEMATVASSSSLEK, which was the only one identifying the POTE ankyrin domain family member F protein (ID A5A3E0). It was only reported by DIA-NN in one of three technical replicate analyses (Sample A6_1_2452) but with a high peptide q-value of 7*10^−3^. To assure that the protein is really present, we investigated the raw data signal of this peptide in AlphaViz (Fig. 5). For the replicate in which the peptide was identified, the position of the peptide on the MS1 heatmap is in a crowded part of the ion cloud, but the zoomed-in view revealed no interfering peptides (Fig. 5). The XICs and heatmaps for the peptide and fragments (b3-b8 and y3-y8 fragment ion series) confirm the presence of the peptide. Taking into account the information about the detected peptide, such as its retention time, charge, and ion mobility, we were able to retrieve this peptide signal in the other two replicates where this peptide had not been reported. Interestingly, we found high quality signals for this peptide comparable to the first sample in both remaining copies, suggesting that improvements to the software could in the future lead to even higher data completeness.

**Fig. 5.**
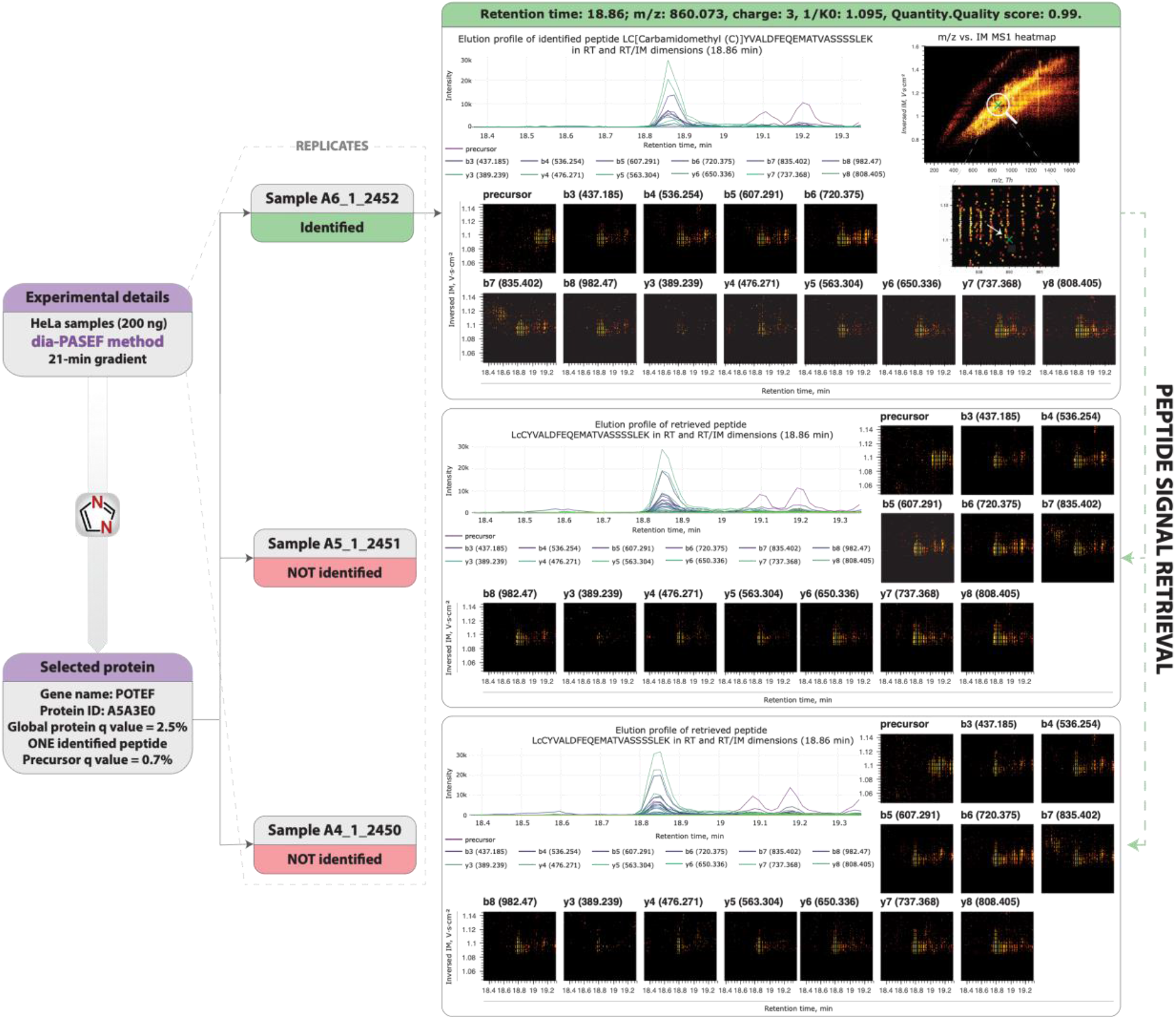
Validation of peptide signal presence in all replicates of an experiment. *Left part, Experimental details*. Three replicates of a HeLa analysis on a timsTOF pro instrument analyzed by DIA-NN (PXD017703) (26, 27). The POTE ankyrin domain family member F protein (ID A5A3E0) with only a single cystein-carboxylated peptide LCYVALDFEQEMATVASSSSLEK identified in one replicate (Sample A6_1_2452) was selected for further investigation. *Right part, Extracted peptide signals*. Raw data extracted for the identified peptide (top). Based on the known peptide sequence, the raw signal of this peptide was successfully extracted from the two remaining samples in which the peptide was not initially identified.

### Data quality assessment and discovery of EGF signaling events using AlphaViz

Studies on post-translational modifications (PTMs) by their nature rely on the identification and quantification of single peptides and are especially affected by poor peak qualities and missing values, the two major challenges AlphaViz tries to overcome. We investigated signalling events activated along the well-studied epidermal growth factor (EGF) signaling pathway. Binding of EGF to its receptor EGFR induces a signalling cascade mediated by phosphorylation leading to cellular proliferation, differentiation and survival (43). We used a recently published dataset where Hela cells were stimulated with EGF or left untreated, acquired in three replicates each on a timsTOF pro instrument with a 21-min gradient in dia-PASEF mode and analyzed with DIA-NN (Experimental Procedures) (28). When filtered for 100% valid values in each condition, DIA-NN detected 1,403 phosphosites as significantly upregulated, of which 56 were localized on proteins known to be part of the EGFR signaling pathway (according to Gene Ontology Biological Process (GOBP) (44). To evaluate the data quality of regulated phosphosites, we picked significantly upregulated phosphosites with DIA-NN scores > 0.7 (FDR < 0.05). The majority of regulated phosphosites with higher DIA-NN scores showed well correlating elution profiles of precursor and fragment ions, for example the peptide carrying the phosphorylation on S642 of RAF1 (Fig. 6A, green in Fig. 6B). However, others demonstrated poor data quality (red in Fig. 6A). This also affected phosphorylation on proteins known to be associated with the EGFR signaling pathway that are presumably correct. For example, the CBL (Casitas B-lineage Lymphoma) protein, an E3 ligase known to ubiquitylate EGFR showed poor quality elution profiles for its peptides phosphorylated on S619 and S667 despite a maximum DIA-NN score between replicates of 0.92 and 0.94 respectively (red in Fig. 6B) (45). Specifically, AlphaViz only retrieved an elution peak for the precursor but none of the expected fragments co-eluted. This was the case for a number of peptides in the EGFR pathway (red in Fig. 6A). We assume that the neural network in DIA-NN scored the presence of the peptide in these cases mainly based on the precursor. While this may be justified in these cases, it would be problematic without supporting biological a priori information. We hope that this observation will initiate improvement to software tools – for instance it could be reported that matching was only based on MS1 level. In any case, we recommend to employ AlphaViz for data quality checks before extensive follow-up experiments.

**Fig. 6.**
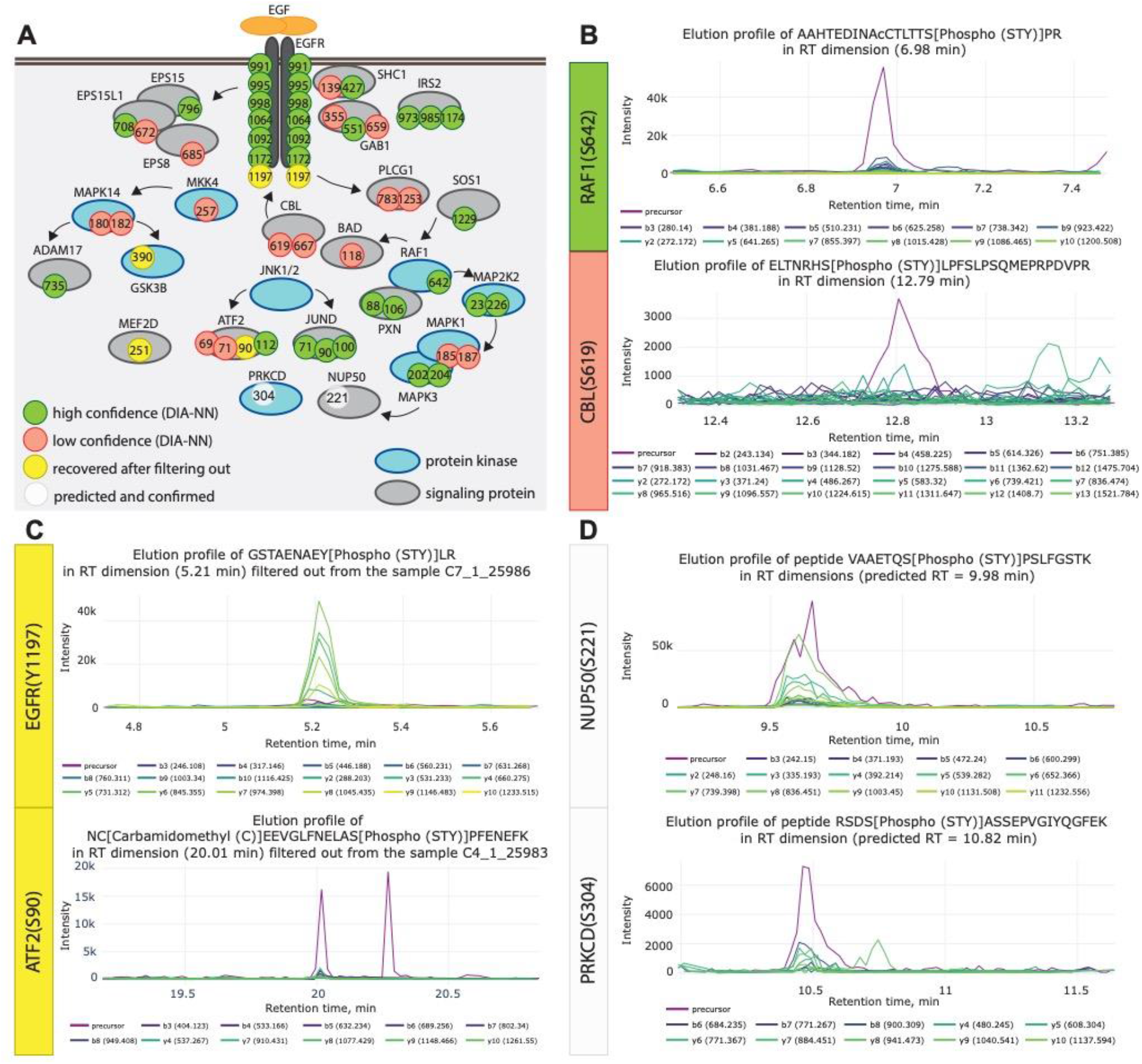
Investigation of EGF-induced phosphorylation events in AlphaViz. *A, Scheme of significantly upregulated phosphosites investigated in AlphaViz*. Based on visual inspection, we divided the reported phosphosites into four main groups: high confidence (green), low confidence (red), recovered after filtering (yellow), and predicted and confirmed (white). *B*, Elution profiles of two phosphorylation sites with high (S642 of RAF1) and low (S619 of CBL) confidence. *C*, Elution profiles of two phosphorylation sites (Y1197 of EGFR and S90 of ATF2) recovered after filtering out. *D*, Elution profiles of two phosphorylation sites (S221 of NUP50 and S304 of PRKCD) not reported by DIA-NN but found in the data using predict mode.

A second challenge are missing values, especially in PTM studies. Although this problem is much reduced in DIA compared to DDA data, it still occurs frequently especially as sample size grows (39, 46–48). This is due to the complexity of spectra, the low abundance of modified peptides and the technical variability. Unfortunately, it has been impossible or extremely laborious to manually check raw data for specific spectra or elution groups of modified peptides that were not reported by the proteomics workflows. The predict mode in AlphaViz addresses this issue. It only requires the peptide sequence, peptide charge and type and localization of its modification to predict its retention time, ion mobility and fragment intensities.

We first investigated functional phosphorylation events on proteins involved in EGF signalling that showed increased intensities upon EGF treatment, but were lost due to filtering the dataset for 100 % valid values in at least one condition. In most of these cases, the data quality of phosphopeptides in replicates where the DIA software did not report intensities was comparable to the respective replicates with reported intensities. This affected phosphorylation events on proteins along the whole EGF signalling pathway starting with the EGFR receptor itself (Y1197), kinases regulating downstream signalling like GSK3B (T390) and phosphorylation events activating transcription factors like MEF2D (S251) and ATF2 (S90) (Fig. 6C, Supplementary Fig. S4). These examples are clearly false negatives of the computational pipeline and they prove the potential of our tool to recover biologically correct regulatory sites. In the case of novel sites, AlphaViz could have prevented them from being discarded because of data incompleteness.

Besides these reported regulatory phosphosites, the predict mode also provides the possibility to look for phosphorylation events in the raw data that have not been identified by the proteomics software at all. In these cases, AlphaViz uses the peptide sequence, PTM localization and charge state of modified peptides of interest to retrieve the corresponding locations in the raw data. To illustrate, in the EGF dataset, we would have expected increased phosphorylation of the nuclear pore complex protein 50 (NUP50) on position S221, which is mediated by the extracellular signal-regulated kinases (ERK) downstream of the EGF receptor (49), but no such peptide was reported. Remarkably, elution group profiles at the predicted retention time and ion mobility were of good quality and confirmed the presence of this phoshopeptide in the sample (Fig. 6D). This was also the case in a second example, relating to EGF-induced activation of the protein kinase C delta (PRKCD), a kinase regulating cell-adhesion upon EGF stimulation, which leads to its autophosphorylation at S304 (50, 51). Hence, the predict mode allows us to efficiently investigate specific signalling events of interest that were missed by MS software tools through direct inspection of the raw data.

## DISCUSSION

Increasingly automated and capable proteomics processing workflows provide researchers with an easy way to summarize the results of proteomics experiments with statistical confidence measures. However, expert evaluation of individual peptides and proteins is lost along the way. To remedy this, we have developed AlphaViz, a Python-based software package for easy visual validation of critical identifications. Like other members of the AlphaPept ecosystem, it adheres to modern and robust software development principles and it is available to community as a Python module and the GUI for end users.

Future implementations of AlphaViz will include the support for more MS platforms, especially Orbitrap instruments, as well as the integration of results from other used proteomics software packages. Furthermore, due to the well-documented and tested open-source code, AlphaViz is easily extendable by bioinformaticians who want to integrate the latest cutting-edge ideas, as already demonstrated by AlphaPept and AlphaTims (13, 16). Additionally, directly linking protein candidates from fully automated downstream analysis packages like the clinical knowledge graph will further strengthen the link between raw data and biological insight (14). Since the visualization capabilities of AlphaViz are only limited by data structure, it can also be used for the in-depth inspection of lipidomics and metabolomics data.

Here we have demonstrated how AlphaViz can quickly give the researcher confidence in identified, critical peptides by inspection of the search results with the raw chromatographic, ion mobility, MS1 and MS2 levels. Conversely, visualization strongly suggests that some peptides are likely false positives despite of their high search engine scores. Furthermore, AlphaViz makes use of the revolution in deep learning enabled prediction of experimental peptide properties from the identified amino acid sequence. This feature is the basis for the ‘predict mode’, in which we retrieve the raw data for peptides that were not reported by the automated workflow, but were potentially present in the data. This may allow the rescue of low-level signals that are biologically expected to be present or are present in some but not all replicates. In phosphoproteomics of the EGF signaling pathway, we showed how this can help to validate the presence of reported signal nodes, and to avoid extensive follow-up experiments for novel phosphorylation sites whose raw data make them unlikely to be true.

In conclusion, we believe researchers will profit from the minimal time investment to visually check their critical peptides and proteins, potentially saving the community and themselves from futile follow up work. This is particularly true of very surprising and biologically unexpected results that then fail to be reproduced by the wider community. In this context, journals could encourage or mandate the inclusion of such extra data and visualizations for the critical peptides or peptidoforms that form the basis of the new hypotheses, helping to address the ‘crisis of reproducibility’ (52, 53).

## ABBREVIATIONS

BPI: base peak intensity
CCS: collisional cross section
DDA: data-dependent acquisition
DIA: data-independent acquisition
EGF: epidermal growth factor
FDR: false discovery rate
GOBP: Gene Ontology Biological Process
GUI: graphical user interface
IM: ion mobility
IQR: interquartile range
MS/MS: or MS2 tandem MS
PASEF: parallel accumulation – serial fragmentation
PEP: posterior error probability
PTM: post-translational modification
PyPI: Python Package Index
RT: retention time
TIC: total ion current
TIMS: trapped ion mobility spectrometry
TOF: time-of-flight
XIC: extracted ion chromatogram

## ACKNOWLEDGEMENTS

We thank our colleagues in the Department of Proteomics and Signal Transduction (Max Planck Institute of Biochemistry). We are grateful to Stephan Uebel and Stephan Pettera from the core facility and to Özge Karayel for providing experimental data on phosphoisomers. We are thankful to Maria Wahle and Corazon Ericka Mae Itang for testing and providing critical feedback on AlphaViz, and to Medini Steger for great help in editing the manuscript and user manual.

## FUNDING AND ADDITIONAL INFORMATION

This study was supported by the Bavarian State Ministry of Health and Care through the research project DigiMed Bayern (www.digimed-bayern.de), by the Max-Planck Society for Advancement of Science and by the Deutsche Forschungsgemeinschaft (DFG) project ‘Chemical proteomics inside us’ (grant 412136960). M.T.S is supported financially by the Novo Nordisk Foundation (Grant agreement NNF14CC0001).

## DATA AVAILABILITY

The source code and user guide are available under Apache 2.0 license and can be found on GitHub at https://github.com/MannLabs/alphaviz. The mass spectrometry proteomics data have been deposited to the ProteomeXchange Consortium *via* the PRIDE partner repository (54) with the dataset identifier PXD034223.

## AUTHOR CONTRIBUTIONS

E.V and M.M. conceptualized and designed the study; P.S. performed the experiments; E.V. implemented the Python code and GUI; S.W., W-F.Z., and M.T.S. reviewed and contributed the code; E.V., S.W., P.S., M.C.T., and M.M. analyzed the data; S.W., A.-D.B., P.S., and M.T. provided valuable ideas for the concept and visualization in AlphaViz; E.V., S.W., P.S., M.C.T., A.-D.B., and M.M. wrote the manuscript; S.W. and M.M. coordinated and supervised the study.

## SUPPLEMENTAL DATA

This article contains supplemental data.

