## Supplemental files for "AlphaViz: Visualization and validation of critical proteomics data directly at the raw data level"

### SUPPLEMENTARY FIGURE LEGENDS

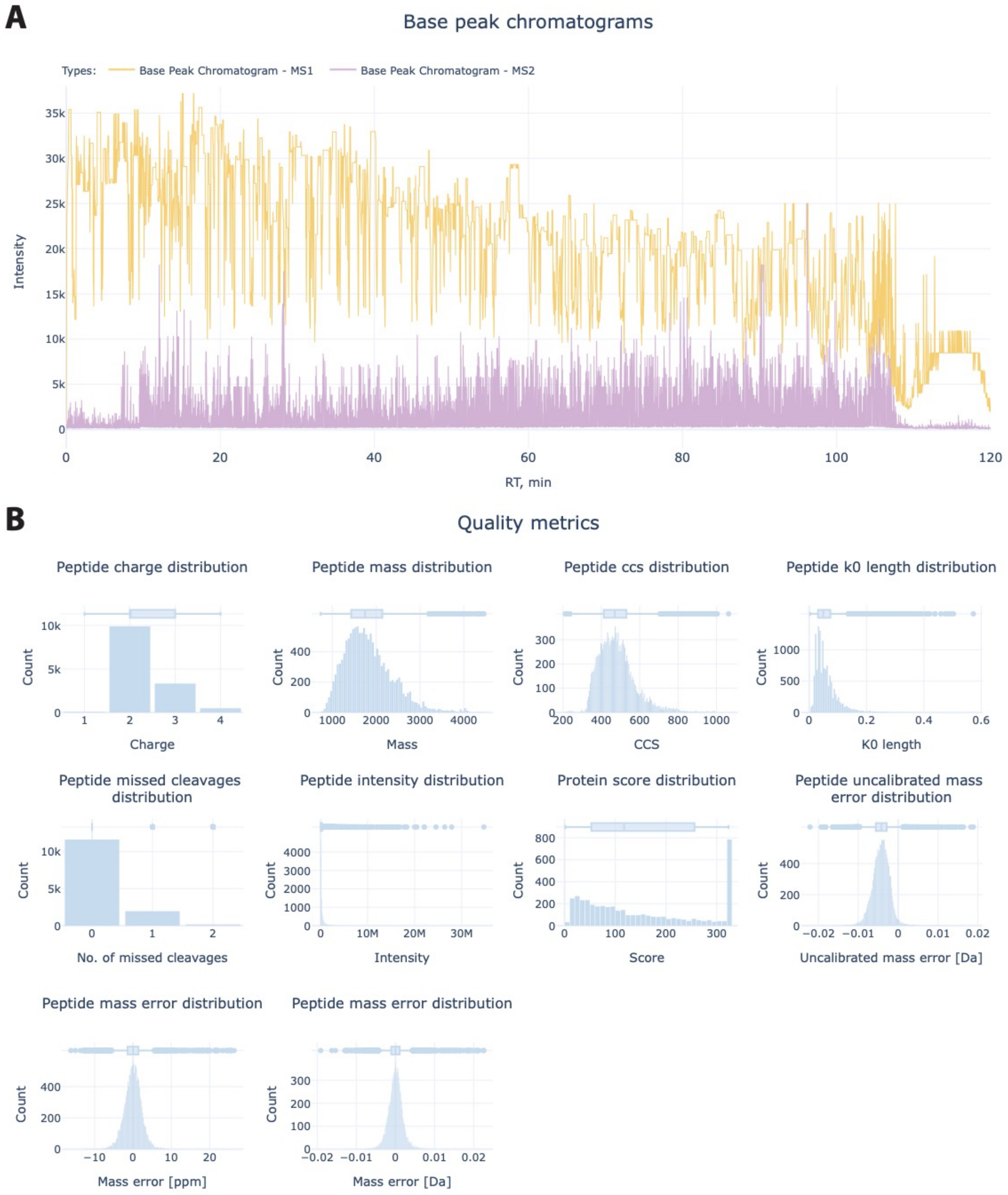


*Supplementary Figure S1.*

A) Base peak intensity chromatograms of MS1 and MS2 data from 120-min HeLa sample, acquired with dda-PASEF method (PXD010012) (25).

B) Ten additional quality metrics available in AlphaViz for DDA data analyzed by MaxQuant.

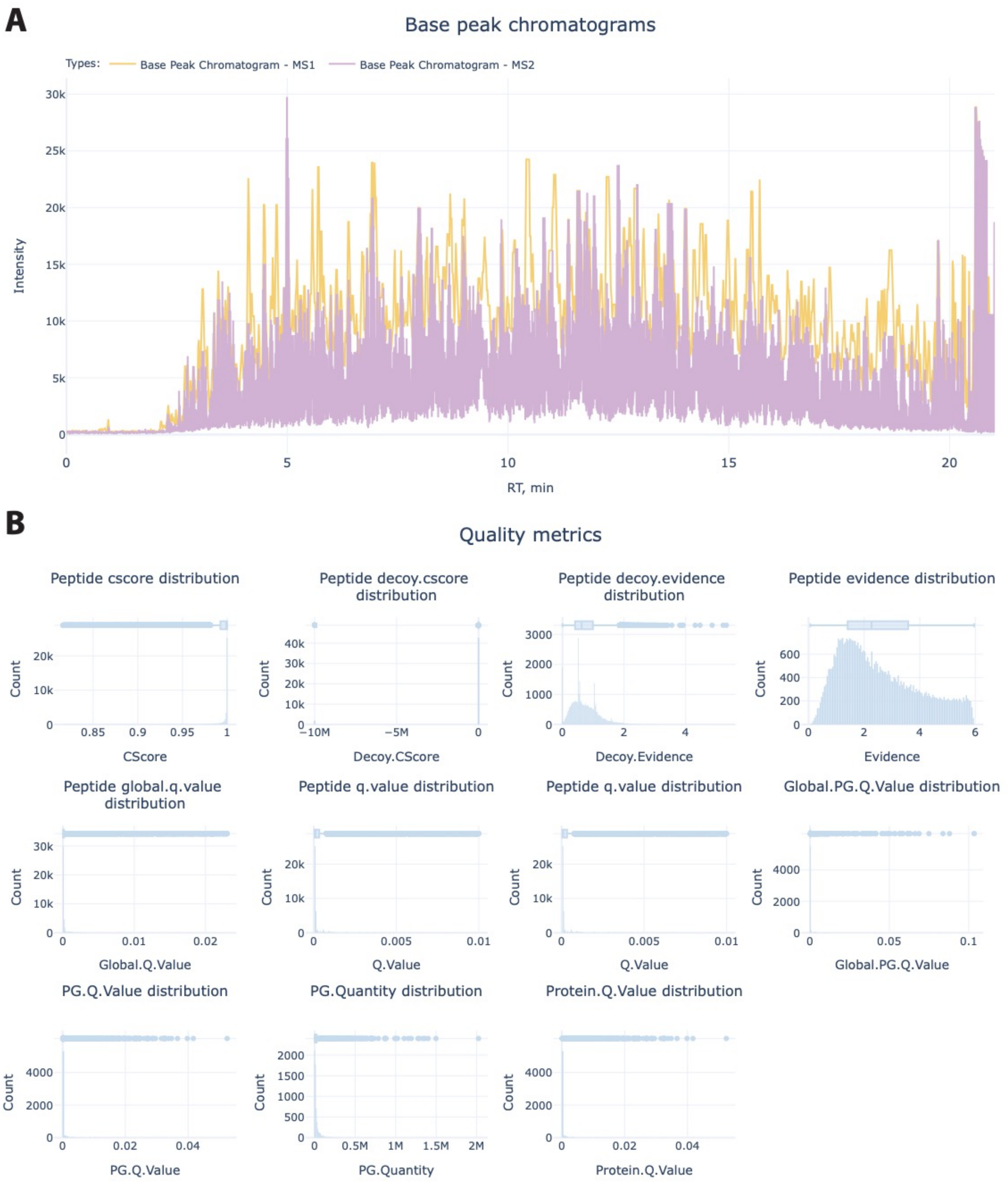


*Supplementary Figure S2.*

A) Base peak intensity chromatograms of MS1 and MS2 data from 21-min HeLa sample, acquired with diaPASEF method (PXD017703) (26).

B) Eleven additional quality metrics available in AlphaViz for DIA data analyzed by DIA-NN.

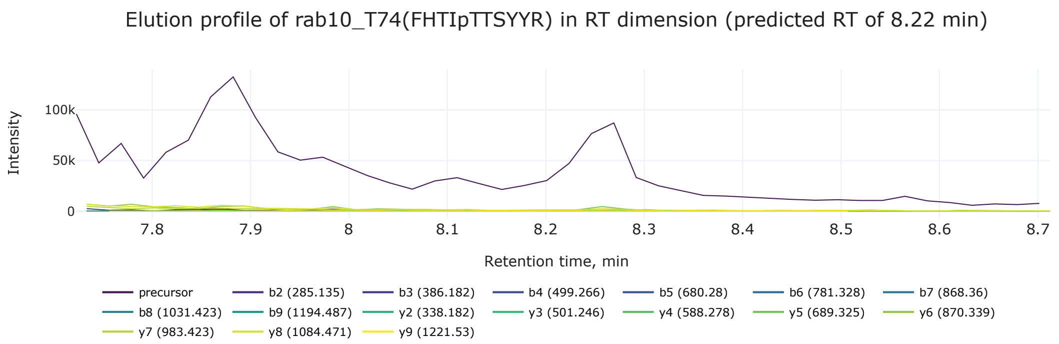


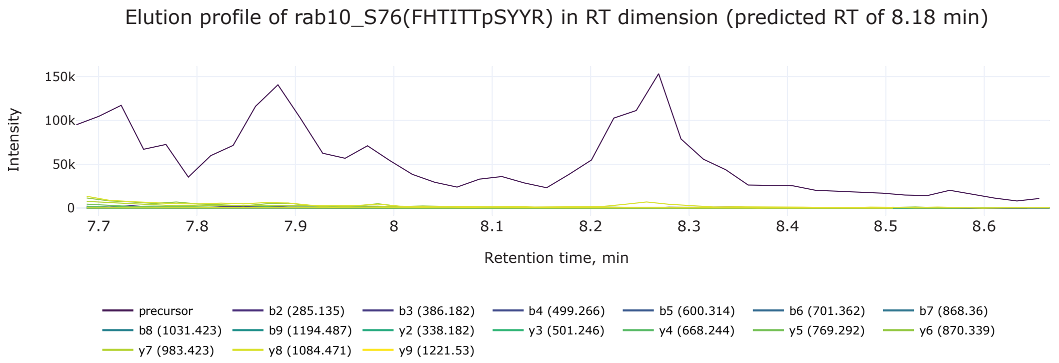


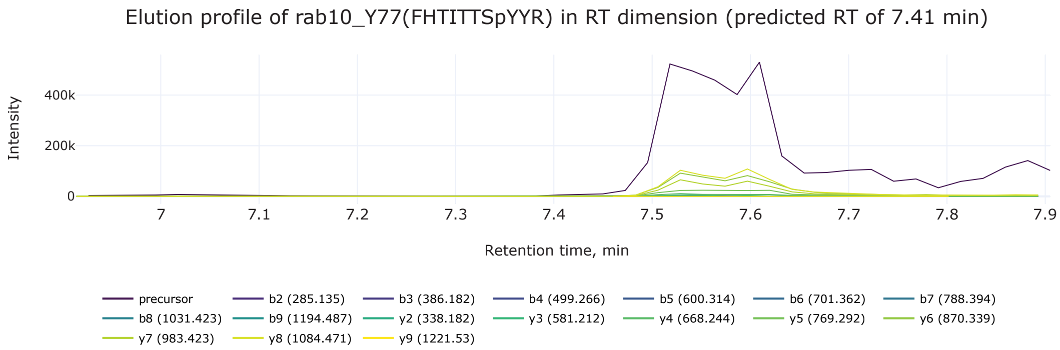


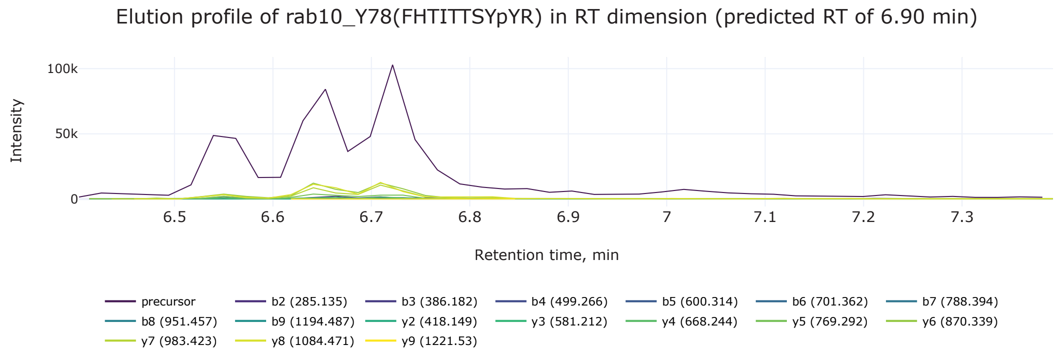


*Supplementary Figure S3.* Elution profiles of the remaining synthetic phosphoisomers of Rab10 peptide in DIA raw data in retention time (RT) dimension.

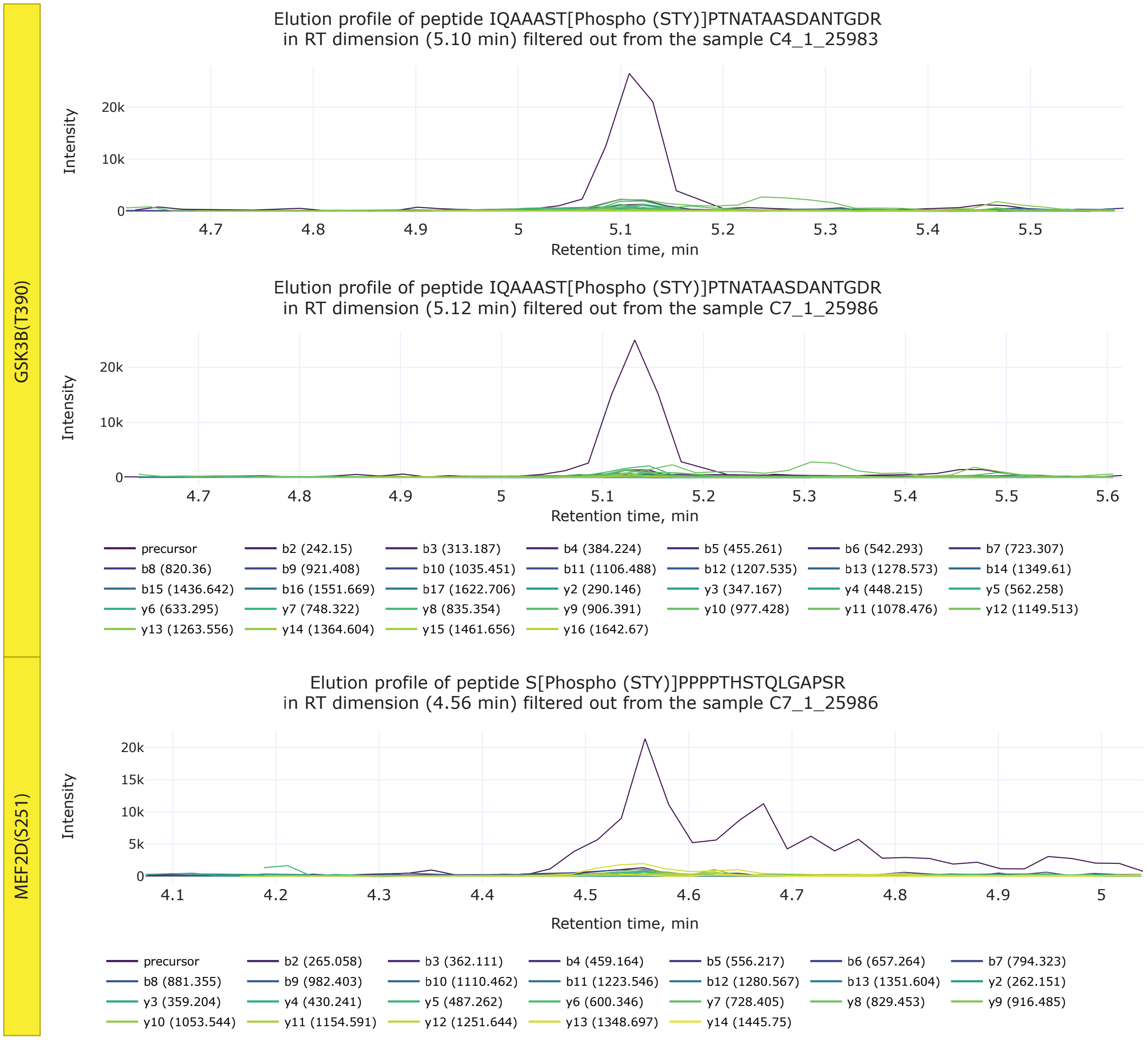


*Supplementary Figure S4.* Elution profiles of two phosphorylation sites (T390 of GSK3B and S251 of MEF2D) in retention time dimension recovered after filtering out.
